# Rare Variable *M. tuberculosis* Antigens induce predominant Th17 responses in human infection

**DOI:** 10.1101/2024.03.05.583634

**Authors:** Paul Ogongo, Liya Wassie, Anthony Tran, Devin Columbus, Lisa Sharling, Gregory Ouma, Samuel Gurrion Ouma, Kidist Bobosha, Cecilia S. Lindestam Arlehamn, Neel R. Gandhi, Sara C. Auld, Jyothi Rengarajan, Cheryl L. Day, John D. Altman, Henry M. Blumberg, Joel D. Ernst, the TBRU ASTRa Study Group

## Abstract

CD4 T cells are essential for immunity to *M. tuberculosis* (*Mtb*), and emerging evidence indicates that IL-17-producing Th17 cells contribute to immunity to *Mtb*. While identifying protective T cell effector functions is important for TB vaccine design, T cell antigen specificity is also likely to be important. To identify antigens that induce protective immunity, we reasoned that as in other pathogens, effective immune recognition drives sequence diversity in individual *Mtb* antigens. We previously identified *Mtb* genes under evolutionary diversifying selection pressure whose products we term Rare Variable Mtb Antigens (RVMA). Here, in two distinct human cohorts with recent exposure to TB, we found that RVMA preferentially induce CD4 T cells that express RoRγt and produce IL-17, in contrast to ‘classical’ *Mtb* antigens that induce T cells that produce IFNγ. Our results suggest that RVMA can be valuable antigens in vaccines for those already infected with *Mtb* to amplify existing antigen-specific Th17 responses to prevent TB disease.

## Introduction

After infection with *Mycobacterium tuberculosis (Mtb)*, the majority of individuals remain well and do not develop active tuberculosis (TB) disease, suggesting that some human immune responses can control the infection. Evidence from people with HIV (Barnes et al. 1991; Sonnenberg et al. 2005; Lawn et al. 2009) and animal models (Scanga et al. 2000; Leveton et al. 1989; Flory et al. 1992; Mogues et al. 2001) have revealed that CD4 T cells are essential for immune control of *Mtb*, but the properties of CD4 T cells that contribute to immune control have not been fully defined. While considerable effort has been focused on identifying T cell effector functions associated with protection, less attention has been given to antigen specificity of T cells as a determinant of immune control, despite the *Mtb* genome encoding ∼4000 potentially antigenic proteins (Cole et al. 1998).

Antigen and epitope specificity are critical determinants of protective immunity in other infectious diseases. Antibodies to pathogens such as Dengue Virus (DV) (de Alwis et al. 2014; Katzelnick et al. 2017), Respiratory Syncytial Virus (RSV) (Polack et al. 2002; Delgado et al. 2009), and *Neisseria meningitidis* (Goldschneider et al.1969) can be associated with protection or not, or even with more severe disease, depending on the antigen and epitope specificity. For infectious diseases in which T cell responses play predominant roles in protection, there is less knowledge of the importance of distinct antigen targets. However, in HIV, there is evidence that CD4 and CD8 T cells that recognize the env protein are associated with poorer control while T cells that recognize gag are associated with better control of viremia (Kiepiela et al. 2007; Ranasinghe et al. 2012). A recent report emphasized the relevance of identifying distinct *Mtb* antigen-specific T cell repertoires as it identified T cell antigen receptor (TCR) clonotypes that are associated with control (maintained latent TB) or progression to active TB disease (Musvosvi et al. 2023).

A characteristic feature of host-pathogen relationships is the coevolutionary arms race in which antigens that induce host-protective (and therefore pathogen-detrimental) immune responses are driven to escape recognition through diversifying evolutionary selection and antigenic variation. Antigenic variation to escape immune recognition has been observed for diverse pathogens, including viruses like HIV (Crawford et al. 2009), HCV (Wang et al. 2010) and influenza (Rimmelzwaan et al. 2009); bacteria including *Streptococcus pneumoniae* (Croucher et al. 2011) and *Neisseria gonorrhoeae* (Seifert et al. 1994); parasites such as *Trypanosoma brucei* (Gkeka et al. 2023) and *Plasmodium falciparum* (Schneider et al. 2023; Chew et al. 2022); and even tumor cells (Matsushita et al. 2012; Marty Pyke et al. 2018; Marty et al. 2017; Hoyos et al. 2022). In an earlier study, we hypothesized that antigenic targets of protective immunity to *Mtb* could be discovered by studying evolutionary selection pressure through T cell epitope sequence variation in phylogeographically diverse clinical isolates. We made the unexpected discovery that the great majority of experimentally verified human T cell epitopes in *Mtb* were perfectly conserved, and exhibited no evidence of antigenic variation (Comas et al. 2010). This result raised the question whether there were undiscovered antigens and epitopes under diversifying selection pressure from human T cell recognition. In a subsequent study, we used comparative genomics to discover *Mtb* genes that exhibit evidence of the strongest diversifying evolutionary selection presssure. We determined that the proteins encoded by those genes are antigenic and that the observed amino acid substitutions impact human T cell responses (Coscolla et al. 2015). We termed these antigenic proteins the Rare Variable *Mtb* Antigens (RVMA).

In the present study, we tested the hypothesis that RVMA express functions distinct from those that recognize classical immunodominant *Mtb* antigens. We first studied 60 distinct *Mtb* antigens (Whatney et al. 2018) using whole blood ELISA, and found that immunodominant ’classical’ *Mtb* antigens elicit higher frequency (% of samples with a measurable response) and magnitude (amount of cytokine, or % of T cells responding) IFNγ responses compared with other *Mtb* antigens, including RVMA. Focusing on four RVMA and four classical antigens from the 60-antigen set, we used intracellular cytokine staining (ICS), to discover that CD4 T cells that recognize RVMA predominantly produce IL-17 and express RORγT, while confirming that classical *Mtb* antigens drive Th1 responses characterized by high IFNγ production and high expression of T-bet and CXCR3. These findings reveal previously unknown skewing of Th17 responses to the rare *Mtb* antigens that are characterized by diversifying evolutionary selection pressure and suggest that TB vaccine strategies will benefit from including antigens that induce T cell responses beyond those of Th1 cells.

## Materials and methods

### Study participants and sample collection

#### Cohort 1

Household contacts (HHCs) of newly diagnosed active pulmonary TB cases were referred to the Kenya Medical Research Institute (KEMRI) Clinical Research Center in Kisumu, Kenya, and their demographic and medical history data were collected. Active pulmonary TB (index) cases with drug-susceptible TB were symptomatic individuals with acid-fast bacilli (AFB) sputum smear positive or a positive GeneXpert MTB/RIF (Cepheid, Sunnyvale, California) result and a positive culture for *Mtb* growth identified at community health clinics in Kisumu, Western Kenya. HHCs were persons who shared the same home residence as the index case for ≥5 nights during the 30 days prior to the date of TB diagnosis of the index case, and were enrolled no more than 3 months (mean: 18 days; range: 1-77 days) after the index case began TB treatment. All participants provided written informed consent to join the study and were recruited from two community-based health clinics located in Kisumu City and Kombewa, Kisumu County. All enrolled individuals met the following inclusion criteria: ≥ 13 years of age at the time of enrollment, positive QuantiFERON TB Gold in Tube (QFT) result, seronegative for HIV antibodies, no previous history of diagnosis or treatment for active TB disease or LTBI, normal chest X-ray, and not pregnant. All participants were presumed to be BCG vaccinated due to the Kenyan policy of BCG vaccination at birth and high BCG coverage rates throughout Kenya. All participants gave written informed consent for the study, which was approved by the KEMRI/CDC Scientific and Ethics Review Unit and the Institutional Review Board at Emory University, USA.

#### Cohort 2

Enrollment criteria and procedures for this cohort were the same as for Cohort 1, with the difference that potential participants (index cases and HHCs) were identified through public health surveillance at community health facilities in Addis Ababa, Ethiopia. All participants provided written informed consent to join the study, which was approved by the Institutional Review Board at Emory University, the AHRI/ALERT Ethics Review Committee, and the Ethiopian National Research Ethics Review Committee.

At both study locations, blood samples were collected from participants in sodium heparin or lithium heparin Vacutainer CPT Mononuclear Cell Preparation Tubes (BD Biosciences or Greiner Bio-One). PBMC were isolated by density centrifugation, rested in complete media (RPMI 1640 containing L-glutamine supplemented with 10% heat-inactivated fetal bovine serum (FBS), 1% PenStrep, 1% Hepes) before counting, and stored in liquid nitrogen (LN2) until use. PBMC isolation was initiated <2 hours after the blood was drawn. Isolated PBMC were cryopreserved in 90% heat-inactivated fetal calf serum/10% DMSO, and kept in LN2 until they were thawed for study at the UCSF laboratory.

### Mycobacterium tuberculosis antigens

Sixty distinct *Mtb* antigens were synthesized as peptide pools of 18 amino acids overlapping by 11 as previously described (Whatney et al. 2018). These antigens are derived from proteins in different bacterial functional categories including cell wall and cell processes; intermediary metabolism and respiration; virulence, detoxification, and adaptation; information pathways; lipid metabolism, and conserved hypotheticals. Additionally, the antigens are derived from different fractions of the bacteria including membrane, secreted, cytoplasm, cell wall, and predicted membrane/secreted proteins. Several of the antigens were initially identified by their exhibiting evidence of antigenic variation and diversifying evolutionary selection pressure. For those antigens, the overlapping peptide pools included all of the known sequence variants identified in (Coscolla et al. 2015).

### Whole blood Response Spectrum Assay (RSA)

Heparinized whole blood was processed within 2 hours of collection as previously described (Whatney et al. 2018). Blood was diluted at 1:4 with media consisting of RPMI 1640 medium supplemented with 2 mM L-glutamine, 100 U/ml penicillin, and 100 mg/ml streptomycin. Diluted blood (100μl) was added to each well of a sterile 96-well round-bottom, tissue culture–treated plate (Corning) containing 100μl of RSA medium alone (negative control), RSA medium with individual *Mtb* antigen prepared as peptide pools at 1μg/ml final concentration, or RSA medium with PHA (positive control) to make a final volume of 200μl per well. Plates were incubated in a 37°C incubator with 5% CO2 for 7 days. On day 7, plates were centrifuged at 900 x g for 5 min, and 150μl of supernatant was removed from each well and transferred to a V-bottom 96-well plate (Corning). Plasma supernatants were stored at -80°C until use for ELISAs. The assays were performed in laboratories at KEMRI, in Kisumu, Kenya.

### IFNγ ELISA

IFNγ in supernatants from the whole blood RSA was determined by ELISA according to the manufacturer’s instructions (Human IFNγ Uncoated ELISA kit; Invitrogen). 50μl of supernatant was diluted with 50μl of assay diluent for use in the IFNγ ELISA. ELISA plates were read at 450 nm (Synergy H1 Microplate reader, Biotek), and data collected using Gen5 software (Biotek).

### PBMC antigen stimulation and intracellular cytokine staining

Cryopreserved PBMCs were thawed in a 37°C water bath until there were ice-balls in the cryotubes, then quickly transferred into pre-warmed R10 media (RPMI 1640 containing L-glutamine supplemented with 10% FBS, 1%PenStrep and 1% Hepes). Cells were centrifuged at 900 x g for 5 minutes at room temperature, the supernatant was discarded, and then cells were resuspended before adding 5ml of warm R10 media and transferred to a 6-well culture plate overnight at 37°C/5% CO2.

Rested cells were transferred to 15ml tubes and live cells were counted using Trypan blue exclusion. 1x10^6^ live cells in 200µL R10 were transferred to each well of a 96-well round bottom culture plate and stimulated with peptide pools (2µg/ml) representing single *Mtb* antigens or SEB as a positive control (1µg/ml). Unstimulated samples were included for each participant as negative controls. Anti-CD28 (1µg/ml) and anti-CD49d (1µg/ml) costimulatory antibodies (both BD Biosciences) were added to each well and the cells were incubated for 2h before adding GolgiStop and GolgiPlug (both from BD Biosciences). The cells were incubated for an additional 18 hours.

After 20 hours of total stimulation, cells were centrifuged at 900 x g for 5 minutes at room temperature and stained with Zombie Aqua Fixable viability kit (Biolegend, 1:1000 diluted in PBS, 100µL) for 20 minutes at room temperature in the dark to enable exclusion of dead cells. 100µL of PBS was added to each well and centrifuged at 900 x g for 5 minutes at room temperature. Cells were washed one more time with 200µL PBS and supernatants were discarded. Next, the cells were resuspended in 50µL of surface antibody cocktail diluted in Brilliant Violet buffer (BD Biosciences) for 20 minutes in the dark. The surface antibody cocktail consisted of αCD3 Brilliant Violet 421 clone UCHT1 (BioLegend) or PE-CF594 clone UCHT1 (BD Biosciences), αCD4 Brilliant Violet 605 clone SK3 (BioLegend), αCD8 Brilliant Violet 650 clone RPA-T8 (BioLegend) or Brilliant Violet 570 clone RPA-T8 (BioLegend), αCD45RA Pacific Blue clone HI100 (BioLegend), αCCR7 Brilliant Violet 785 clone G043H7 (BioLegend), αCD95 PerCP/Cyanine5.5 clone DX2 (BioLegend) and αCD27 FITC clone O323 (BioLegend). 150µL of PBS was added into each well and cells were centrifuged at 900 x g for 5 minutes at room temperature, the supernatant was discarded, and cells were washed again with 200µL PBS. Cells were then permeabilized and fixed using BD Cytofix/Cytoperm kit (BD Biosciences) according to the manufacturer’s instructions for 20 minutes at 4°C protected from light. Cells were washed twice (900 x g for 5 minutes each) with 1X BD Perm/Wash solution. Cells were resuspended in 50µL intracellular antibody cocktail diluted in 1X BD Perm/Wash for 20 minutes at room temperature. The cytokine staining cocktail consisted of αTNF Alexa Fluor 700 clone MAb11 (BD Biosciences), αIFNγ APC/Cyanine7 clone 4S.B3 or PE/Cyanine7 clone 4S.B3 (both BioLegend), αGM-CSF APC clone MP1-22E9 (BioLegend) and αIL-17 PE clone BL168 (Biolegend). After intracellular staining, cells were washed twice with 1X BD Perm/Wash solution. Cells were then fixed in 2% PFA and acquired with an LSR-II cytometer (BD Biosciences).

### T cell activation induced marker (AIM) assay

Cryopreserved PBMCs were thawed, rested overnight, and 1x10^6^ live cells in 200µL R10 were transferred to each well of a 96-well round bottom culture plate. Costimulatory antibodies anti-CD28 (1µg/ml) and anti-CD49d (1µg/ml) together with anti-CD40 (1µg/ml) (Milteny Biotec) were added into each well and rested for 15 minutes in 37°C/5%CO2 incubator. Next, peptide pools representing single *Mtb* antigens at a final concentration of 2µg/ml and SEB positive control (1µg/ml) were used to stimulate the cells for a total of 20 hours. Wells with no stimulus were included as negative controls for each participant in each assay. At the end of stimulation, cells were centrifuged at 900 x g for 5 minutes at room temperature and surface stained with Live/Dead Fixable Blue Dead Cell stain kit (Invitrogen) for 20 minutes at room temperature in the dark. Cells were washed twice (900 x g for 5 minutes at room temperature) and then 50µL of surface antibody cocktail diluted in Brilliant Violet buffer (BD Biosciences) added and incubated 20 minutes at room temperature in the dark. The surface antibody cocktail consisted of αCD3 Brilliant Violet 510 clone UCHT1 (BioLegend), αCD4 BUV496 clone SK3 (BD Biosciences), αCD8 Brilliant Violet 570 clone RPA-T8 (BioLegend), αCD154 PE-CF594 clone TRAP1 (BD Biosciences), αCCR6 FITC clone G034E3 (BioLegend) and αCXCR3 Brilliant Violet 650 clone G025H7 (BioLegend). 150µL of PBS was added into each well and cells centrifuged at 900 x g for 5 minutes at room temperature, supernatant discarded, and cells washed again with 200µL PBS. Cells were then permeabilized and fixed using eBioscience FOXP3/Transcription Factor staining kit (Invitrogen) according to the manufacturer’s instructions for 20 minutes at 4°C protected from light. Cells were washed twice with 1X eBioscience Perm diluent, then resuspended in 50µL of intracellular antibody mix of αRORγT Alexa Fluor 647 Clone Q21-559 (BD Biosciences) and αT-bet PE-Dazzle 594 clone 4B10 or PE clone 4B10 (both BioLegend) diluted in eBiosciences perm diluent. After transcription factor staining, cells were washed twice with 1X eBioscience Perm diluent. Cells were fixed in 2% PFA and acquired using a 5-Laser Cytek Aurora Spectral Flow cytometer (Cytek Biosciences).

### Data and statistical analysis

IFNγ ELISA data was analyzed using SoftMax Pro v6.3 software (Molecular Devices). IFNγ release for each antigen was determined by subtracting the mean background IFNγ concentration in six negative control wells lacking antigens for each participant sample. A maximum concentration of quantifiable IFNγ was set at 1000 pg/ml, corresponding to the concentration of the highest standard of recombinant human IFNγ (Human IFNγ Gamma Uncoated ELISA kit, Invitrogen). IFNγ concentrations below the level of detection by the ELISA standard curve were set to 0 pg/ml. Spectral flow fcs files were initially analyzed using SpectroFlo software v3.0 (Cytek Biosciences) for unmixing and autofluorescence correction. Identification of distinct T cell populations from unmixed fcs files (spectral flow data) and compensated fcs files (LSR-II data) was performed using Flowjo v10 (Flowjo LLC). Antigen-specific cytokine-producing CD4 T cell magnitudes are reported after subtraction of the values from unstimulated samples for each participant; results greater than those of unstimulated samples were considered positive, with the limit of detection set at 0.001% of total CD4 T cells. All statistical analyses were performed using GraphPad Prism version 9.0 or 10 (GraphPad Software, Inc). Comparisons of 2 groups were done by a paired or unpaired 2-tailed Student’s t-test (Wilcoxon’s test or Mann-Whitney test, respectively), and p < 0.05 was considered statistically significant. For calculation of the response magnitudes, samples that yielded undetectable results were entered as 0.001% of CD4 T cells.

## Results

### Individual Mtb antigens elicit IFNγ responses that vary in frequency and magnitude

To characterize the frequency and magnitude of responses to 60 individual *Mtb* antigens, we assayed samples from 71 QFT^+^HIV^-^ adults after recent (≤ 3 months) household exposure to active pulmonary TB from western Kenya (Cohort 1). After quality control and subtraction of background, we found considerable diversity in IFNγ secretion by antigen (columns) and by participant (rows) (Fig. 1A). By arranging the antigens by the position of their genes on the *Mtb* chromosome, the results revealed apparent ‘antigenic islands’ and ‘antigenic deserts’ reflected in the frequency (% of samples with responses) and magnitude (amount of IFNγ produced) of responses. The antigenic islands largely comprise known *Mtb* immunodominant antigens, including EsxH (Tb10.4), PE13, PPE18, Ag85B, PPE46, Ag85A, CFP-10, and ESAT-6 (median values, 50-279 pg/ml), consistent with other reports that these antigens induce IFNγ responses. In some samples, responses to these antigens exceeded the upper limit of the IFNγ assay (1,000 pg/ml), contributing to underestimation of the response magnitudes.

**Figure 1:**
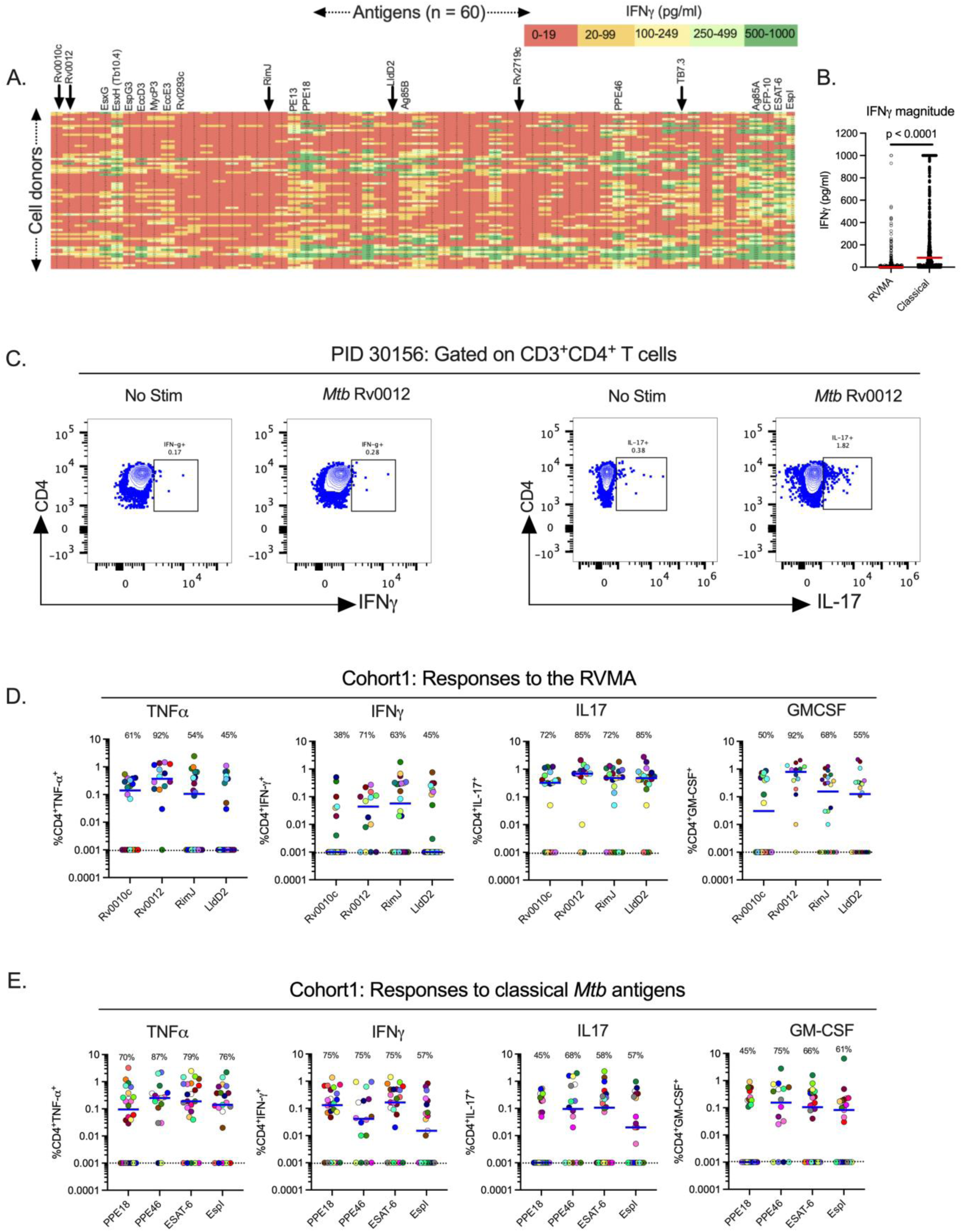
Distinct *Mtb* antigens elicit T cell responses with different functional properties (Cohort 1). **(A)** Whole blood samples from QFT^+^HIV^-^ participants in Cohort 1 (KEMRI, Kisumu, Kenya) were stimulated with 60 individual *Mtb* antigens as overlapping peptides (1μg/ml) for 7 days and supernatants harvested to quantitate IFNγ by ELISA. The antigens are arranged left to right (1-60) according to their positions on the *Mtb* chromosome, and individual participant samples are arranged in rows. Data presented are results after subtraction of the average of six unstimulated wells as the background response. IFNγ levels are color-coded according to magnitude with red for the lowest and green for the highest responses. Responses greater than the highest standard were assigned the value of the highest standard (1000pg/ml). Selected antigens are highlighted at the top of the heatmap with arrows indicating the RVMA, four of which were studied in more detail using flow cytometry. The identity of all of the antigens ordered in the same fashion is available in (Whatney et al. 2018). (**B**) Whole blood IFNγ response magnitude of selected classical antigens (Tb10.4 (EsxH), PE13, PPE18, Ag85B, PPE46, Ag85A, CFP-10, ESAT-6 and EspI) compared to the six RVMA (Rv0010c, Rv0012, RimJ, LldD2, Rv2719c and TB7.3), the red horizontal line represents the median response. Statistical significance was determined by Mann-Whitney test. (**C**) Representative flow cytometry plots comparing CD4 T cells producing IFNg or IL-17 in response to stimulation with one of the RVMA (Rv0012), background response for each cytokine is shown for comparison. **(D and E)** Cryopreserved peripheral blood mononuclear cells (PBMCs) were stimulated with indicated antigens (2 g/ml) for a total of 20 hours in the presence of Golgi Stop and Golgi Plug and costimulatory antibodies anti-CD28 and anti-CD49d; cytokine production by CD4^+^ T cells was determined by intracellular cytokine staining and flow cytometry. Data are values after subtraction of unstimulated cells with values lower than the unstimulated control cells indicated below the dotted line (cut-off of positive response). Percent of participants with a detectable response is shown at the top of each antigen plot. Each symbol is a distinct participant, the horizontal blue line indicates the median cytokine response. Results for RVMA are shown in panel **C**, results for the classical antigens are shown in panel **D**.

In contrast to the findings in (Lindestam Arlehamn et al. 2013), our assay platform and participant population did not show evidence for an antigenic island involving the ESX-3 locus. While EsxH (also known as Tb10.4) induced responses of modest magnitudes (median, 102 pg/ml), the other components of the Esx-3 locus (EsxG, EspG3, EccD3, MycP3, EccE3, and Rv0293c) induced minimal IFNγ responses (medians, 4-14 pg/ml).

Compared with responses to the known immunodominant ’classical’ antigens, we found lower magnitude IFNγ responses (0-19 pg/ml; red color) to other *Mtb* antigens. The antigens that induced low magnitude IFNγ responses include Rv0010c, Rv0012, RimJ and LldD2, which we previously identified as showing evidence of evolutionary diversifying selection (shown in arrows) and termed Rare Variable *Mtb* Antigens (RVMA)(Coscolla et al. 2015). While the overall IFNγ responses to RVMA were low, some participants had higher magnitude IFNγ responses (> 100 pg/ml) to certain RVMA, although these were still lower than the magnitude of responses to the classical antigens (Fig. 1B).

We also observed variation in responses to each of the 60 antigens at the level of individual participants (Fig. 1A, rows), emphasizing that responses to a single *Mtb* antigen are not representative of the response to other *Mtb* antigens within an individual. Thus, we confirmed the diversity in responses to *Mtb* infection in humans in agreement with a previous study (Coppola et al. 2016) and found that classical *Mtb* antigens elicit a robust IFNγ response in whole blood of individuals with LTBI while other antigens, including the RVMA, elicit lower frequency and lower magnitude IFNγ responses.

### Mtb antigens induce diverse cytokine responses that vary by antigen class Response Frequencies

Since RVMA are characterized by evidence of diversifying evolutionary selection and T cell epitope sequence variation, we hypothesized that they induce CD4 T cell responses distinct from those induced by conserved immunodominant *Mtb* antigens. We therefore asked whether RVMA induce CD4 T cells that express cytokines other than IFNγ. For this analysis, we selected 4 RVMA that induced low frequency and magnitude IFNγ responses in the 60-antigen whole blood RSA and compared the responses to 4 ’classical’ *Mtb* antigens that induced high frequency and high magnitude IFNγ responses. We stimulated PBMC from participants in Cohort 1 (Supplemental Table 1) with the individual antigens and quantitated CD4 T cells expressing TNFα, ΙFNγ, IL-17, and/or GM-CSF by intracellular cytokine staining and flow cytometry (Fig 1C, Supplementary Fig 1).

Consistent with the results of the whole blood assay, the 4 RVMA (Rv0010c, Rv0012, RimJ, and LldD2) induced IFNγ-producing CD4 T cells in only a modest fraction of participants (38-71%, depending on the individual antigen). Instead, CD4 T cells responding to individual RVMA produced IL-17 in a high proportion (72-85%) of participants. The frequency of participants whose CD4 T cells produced TNFα or GM-CSF in response to the RVMA varied widely (45-92%, depending on the individual antigen) (Fig 1D and Table 1).

**Table 1.** Frequencies of CD4 T cell cytokine responses, by individual antigens.

|  | **RVMA** |  |  |  | **Classical** |  |  |  |
| --- | --- | --- | --- | --- | --- | --- | --- | --- |
|  | Rv0010c<br>n=18 | Rv0012<br>n=14 | RimJ<br>n=22 | LldD2<br>n=20 | PPE18<br>n=24 | PPE46<br>n=16 | ESAT-6<br>n=24 | EspI<br>n=21 |
| TNF | 61 | 92 | 54 | 45 | 70 | 87 | 79 | 76 |
| IFNγ | 38 | 71 | 63 | 45 | 75 | 75 | 75 | 57 |
| IL-17 | 72 | 85 | 72 | 85 | 45 | 68 | 58 | 57 |
| GM-CSF | 50 | 92 | 68 | 55 | 45 | 75 | 66 | 61 |
| The values shown reflect the percent of participants whose samples yielded detectable responses, defined as >0.001% of CD4 T cells after stimulation with the indicated antigens. |  |  |  |  |  |  |  |  |

At the level of individual antigens, all four RVMA induced IL-17-producing CD4 T cells in >70% of the participants, while other cytokine responses varied widely by antigen. Of the RVMA, Rv0010c induced IFNγ and GM-CSF at the lowest frequencies (38% and 50%, respectively). Notably, Rv0012 induced GM-CSF-producing CD4 T cells in a high frequency (92%) of participants, a rate higher than any of the other RVMA or any of the classical antigens. RimJ induced TNFα in only 54% and induced IFNγ and GM-CSF with frequencies intermediate between those of IL-17 and TNFα. LldD2 induced the lowest frequencies of TNFα-, IFNγ-, or GM-CSF-producing CD4 T cell responses (45%, 45%, and 55%, respectively), but induced IL-17-producing cells in 85% (Fig 1D and Table 1).

For comparison, we studied CD4 T cell production of the same four cytokines in response to 4 classical antigens. This revealed considerable variation in the frequency of individuals that responded with CD4 T cell production of TNFα, ΙFNγ, IL-17, and GM-CSF after stimulation with individual classical antigens (PPE18 (Rv1196), PPE46 (Rv3018c), ESAT-6 (Rv3875) and EspI (Rv3876)) (Fig 1E and Table 1). TNFα responses to individual classical antigens were observed in 70-87% of samples, with the highest frequency TNFα responses to PPE46. IFNγ responses to the 4 classical antigens were observed in 57-75% of samples, with little variation in the frequency of responses to PPE18, PPE46, and ESAT-6, and fewer responses to EspI. By comparison, the 4 classical antigens induced IL-17 (45-68%, depending on the antigen) and GM-CSF (46-65%) in fewer participants.

When we compared the frequency of responders (that is, participants whose CD4 T cells expressed a given cytokine in response to *Mtb* antigen stimulation) by antigen class (i.e., classical vs RVMA), IL-17 responses were more frequent for RVMA than for classical *Mtb* antigens, while TNFα, IFNγ, and GM-CSF responses did not differ significantly (Table 2).

**Table 2.** Frequencies of CD4 T cell cytokine responses, by antigen class.

|  | Median % (interquartile range) n = 4 antigens per category |  | Absolute p (Mann-Whitney) |
| --- | --- | --- | --- |
|  | **RVMA** | **Classical** |  |
| TNF | 58 (47, 84) | 78 (72, 85) | 0.3429 |
| IFNγ | 54 (40, 69) | 75 (62, 75) | 0.0857 |
| IL-17 | 79 (72, 85) | 58 (48, 66) | 0.0286 |
| GM-CSF | 62 (51, 86) | 64 (49, 75) | 0.8857 |
| The values shown reflect the median and 25<sup>th</sup> and 75<sup>th</sup> percentiles of the percent of participants whose samples yielded detectable responses, defined as >0.001% of CD4 T cells after stimulation with the indicated antigens. |  |  |  |

These data indicate that *Mtb* antigens vary in their frequency of induction of distinct CD4 T cell effector cytokines. The data also demonstrate that RVMA induce IL-17-producing CD4 T cells more frequently than IFNγ- or TNFα-producing T cells, and they induce IL-17 responses more frequently than do the classical antigens in recently-exposed household contacts that have controlled *Mtb* infection.

### Response Magnitudes

In addition to variation in the frequency of participants whose CD4 T cells responded to individual antigens with expression of different cytokines, we found variation in the magnitude of the CD4 T cell responses, defined as the percent of CD4 T cells that produced the cytokine of interest in response to individual antigens.

For each of the individual RVMA, the magnitude of IL-17 responses was greater than for TNFα, IFNγ, or GM-CSF, with the exception that Rv0012 induced GM-CSF responses that exceeded those of each of the other three cytokines (Fig 1D and Table 3). For the individual classical antigens, the magnitudes of TNFα and IFNγ responses magnitudes were highest, with the exception of PPE46-induced GM-CSF responses that exceeded the magnitude of IFNγ responses. IL-17 responses were present with the lowest magnitudes for all four of the classical antigens.

**Table 3.** Magnitudes of CD4 T cell cytokine responses (% of CD4 T cells) to individual antigens.

|  | **RVMA** |  |  |  | **Classical** |  |  |  |
| --- | --- | --- | --- | --- | --- | --- | --- | --- |
|  | Rv0010c | Rv0012 | RimJ | LldD2 | PPE18 | PPE46 | ESAT-6 | EspI |
| TNF | 0.14 | 0.37 | 0.11 | 0.001 | 0.1 | 0.25 | 0.19 | 0.14 |
| IFNγ | 0.001 | 0.04 | 0.06 | 0.001 | 0.13 | 0.04 | 0.17 | 0.015 |
| IL-17 | 0.33 | 0.69 | 0.48 | 0.48 | 0.001 | 0.01 | 0.11 | 0.02 |
| GM-CSF | 0.03 | 0.79 | 0.16 | 0.13 | 0.001 | 0.154 | 0.11 | 0.19 |
| Values shown for each cytokine and each antigen are median % of all CD4 T cells that express the specified cytokine. Assays that yielded undetectable levels of the stated cytokine were assigned a value of 0.001%. |  |  |  |  |  |  |  |  |

For the RVMA as a group, the highest magnitude responses were for IL-17-producing CD4 T cells (median, 0.46% of CD4 T cells), while the lowest magnitude responses were for IFNγ-(median 0.01%) (Fig 1D and Table 4). GM-CSF- and TNFα-producing CD4 T cells were found with intermediate frequencies (medians of 0.20 and 0.15% respectively). In contrast, for the classical antigens as a group, the highest magnitude responses were for TNFα-producing CD4 T cells (median, 0.19% of CD4 T cells) with slightly lower magnitude responses for IFNγ (0.08%), IL-17 (0.05%), and GM-CSF (0.09%). When comparing cytokine responses by antigen group, differences were significant for IL-17 (RVMA>classical) and IFNγ (classical>RVMA) (Table 4). These analyses revealed that, consistent with the frequencies of responders, the magnitude of IL-17-producing CD4 T cells is higher for RVMA than for classical antigens.

**Table 4.** Magnitudes of CD4 T cell cytokine responses (% of CD4 T cells), by antigen class.

|  | Median % (interquartile range) n = 4 antigens per category |  | Absolute p (Mann-Whitney) |
| --- | --- | --- | --- |
|  | **RVMA** | **Classical** |  |
| TNF | 0.15 (0.001, 0.41) | 0.19 (0.03, 0.56) | 0.1852 |
| IFNγ | 0.01 (0.001, 0.165) | 0.08 (0.001, 0.33) | 0.0107 |
| IL-17 | 0.46 (0.05, 0.80) | 0.05 (0.001, 0.28) | <0.0001 |
| GM-CSF | 0.20 (0.001, 0.61) | 0.09 (0.001, 0.25) | 0.0735 |
| Values shown for each cytokine and each antigen are median % of all CD4 T cells that express the specified cytokine. For statistical analyses, assays that yielded undetectable levels of the stated cytokine were assigned a value of 0.001%. |  |  |  |

To determine whether the IL-17 bias of CD4 T cell responses to RVMA extends to another population, we used the same experimental procedures to study samples from participants recruited in the same manner as for Cohort 1, in an independent cohort in Addis Ababa, Ethiopia (Cohort 2). The two cohorts were comparable in their distribution of age, sex, body mass index (BMI), and hemoglobin A1c (Supplemental Table 1). The distribution of sex was comparable between the cohorts (p = 0.2611, Fisher’s exact test), and when we considered sex as a biological variable, we found no systematic differences in the results within or between cohorts according to sex. Results of Quantiferon testing differed between Cohort 1 and Cohort 2: the former had significantly higher results for TB Antigen minus Nil, while the latter had higher results for Mitogen minus Nil (Supplemental Table 1).

In Cohort 2, the frequency of cytokine responses to RVMA was lower than for classical antigens; the difference was significant for IFNγ and for IL-17, but not for TNFα or GM-CSF (Fig 2A-B, Supplemental Tables 2 and 3). Likewise, the magnitudes of cytokine-producing CD4 T cells in Cohort 2 were lower in response to RVMA compared with classical antigens; the differences in magnitudes between RVMA and classical antigens were significant for TNFα, IFNγ, and IL-17, but not for GM-CSF (Supplemental Table 4). Because of the lower magnitude responses, the IL-17 response magnitudes to the RVMA did not exceed the IL-17 responses to classical antigens. Despite the overall lower magnitude responses, IL-17 responses were higher than other cytokine responses to RVMA (Fig 2A-B, Supplemental Table 4). When we analyzed IL-17 and IFNγ responses to individual RVMA in Cohort 2, we found that the frequencies of IL-17 responses exceeded those of IFNγ responses for all four of the RVMA (Supplemental Table 5). Likewise, the magnitudes of IL-17 responses exceeded those of IFNγ responses for all four of the RVMA; the differences were significant for Rv0012 and LldD2, though not for Rv0010c or Rim J. Comparison of the cytokine response magnitudes to all of the RVMA together revealed that IL-17 responses were significantly greater than TNFα or IFNγ (p = 0.0378 and 0.0018, respectively), but not GM-CSF (Friedman test with Dunn’s correction for multiple comparisons). Despite the differences in the overall magnitudes of CD4 T cell responses to RVMA in Cohort 2 compared with Cohort 1, the results confirmed the findings in Cohort 1 that RVMA predominantly induce CD4 T cells that produce IL-17.

**Figure 2:**
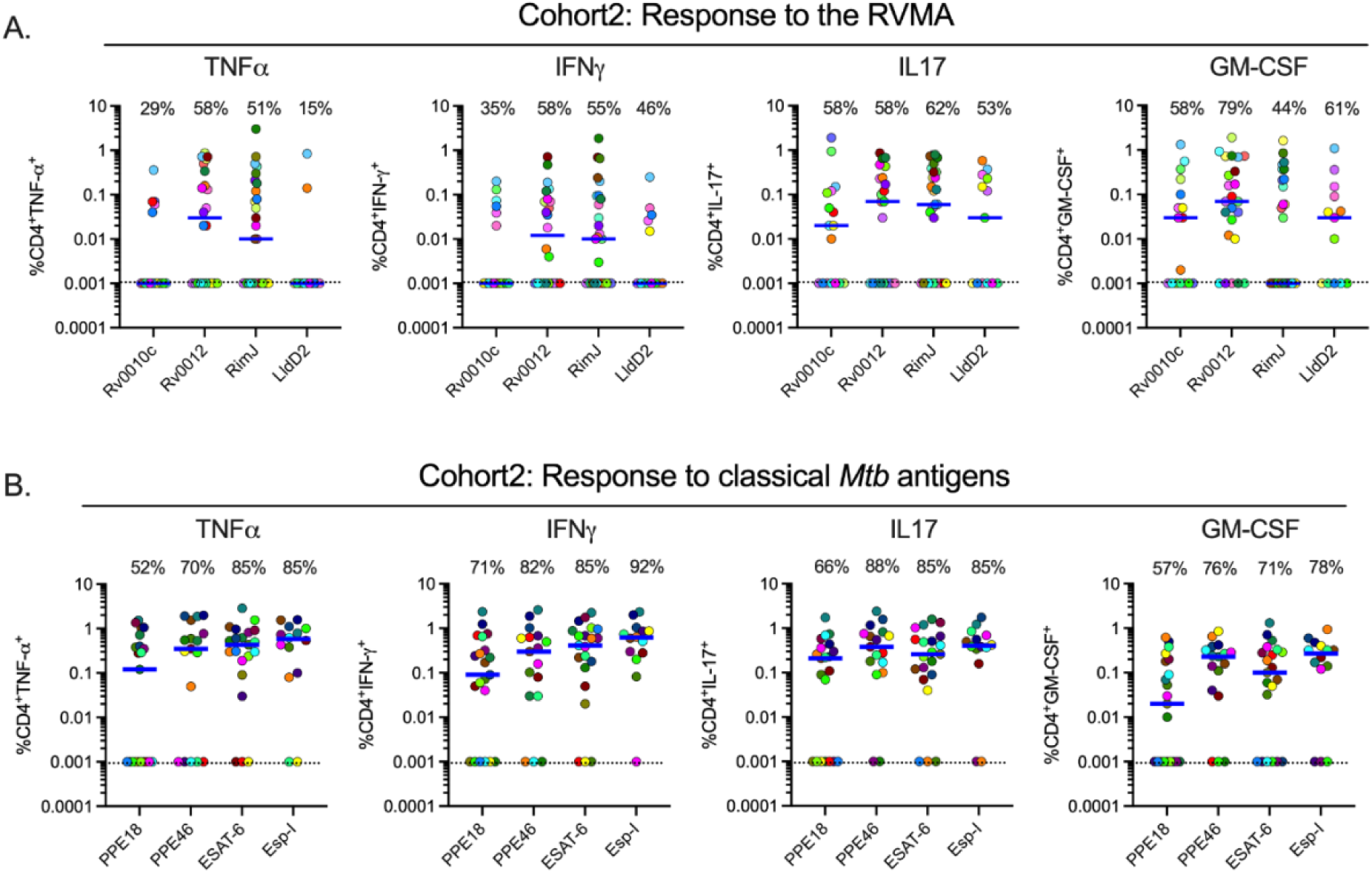
Distinct *Mtb* antigens elicit T cell responses with different functional properties (Cohort 2). Procedures and analyses were as for Figure 1; the samples were obtained from participants in Cohort 2 (AHRI; Addis Ababa, Ethiopia). Results for RVMA are shown in panel **A**; results for classical *Mtb* antigens are shown in panel **B**.

In Cohort 1, RVMA and classical antigens induced similar magnitudes of TNF^+^/IL-17^+^ and IFNγ^+^/IL-17^+^ cells, while RVMA induced lower magnitudes of TNF^+^/IFNγ^+^ cells than did classical antigens. In Cohort 2, RVMA induced lower magnitudes of cells expressing each of the three dual cytokine combinations compared with those induced by classical antigens (Supplementary Figure 2).

### RVMA-responsive CD4 T cells include bona fide Th17 cells

To further investigate CD4 T cells that respond to RVMA, we extended our studies to include additional markers associated with Th17 cells. Since peptide:HLA multimers for the Cohort 1 and Cohort 2 participants’ HLA alleles are not yet available, we used a T cell activation induced marker (AIM) assay (Grifoni et al. 2020; Dan et al. 2016; Barham et al. 2020) based on surface expression of CD154 after brief antigen stimulation in the absence of protein transport inhibitors, and then assayed expression of RORγT and CCR6 (Th17) or T-bet and CXCR3 (Th1) on the activated CD154^+^ cells (Fig. 3A). Stimulation with RVMA induced lower frequencies of CD154^+^ CD4 T cells in PBMC than did classical antigens (Fig. 3B), consistent with immunodominance of classical antigens in this population. When we analyzed lineage-defining transcription factor expression on antigen-responsive (CD4^+^CD154^+^) cells, we found a significantly higher fraction of RORγT^+^T-bet^-^ cells on RVMA-activated cells than on cells activated by classical antigens (Fig. 3C). We also observed a significantly higher fraction of RORγT^-^ T-bet^+^ cells responding to classical antigens than to RVMA. These results are in accord with the cytokine data (Fig. 1 and 2) and indicate that, beyond cytokine production, RVMA-responsive CD4 T cells exhibit other characteristics of Th17 cells while confirming that classical antigen-responsive T cells are typical Th1 cells. We also found a significantly higher frequency of double-negative (RORγT^-^T-bet^-^) antigen-activated cells responding to RVMA than to classical antigens, suggesting potential plasticity or different stages of differentiation of the cells that recognize RVMA. In contrast, there was no difference in double-positive (RORγT^+^T-bet^+^) cells according to antigen category (Fig. 3C).

**Figure 3:**
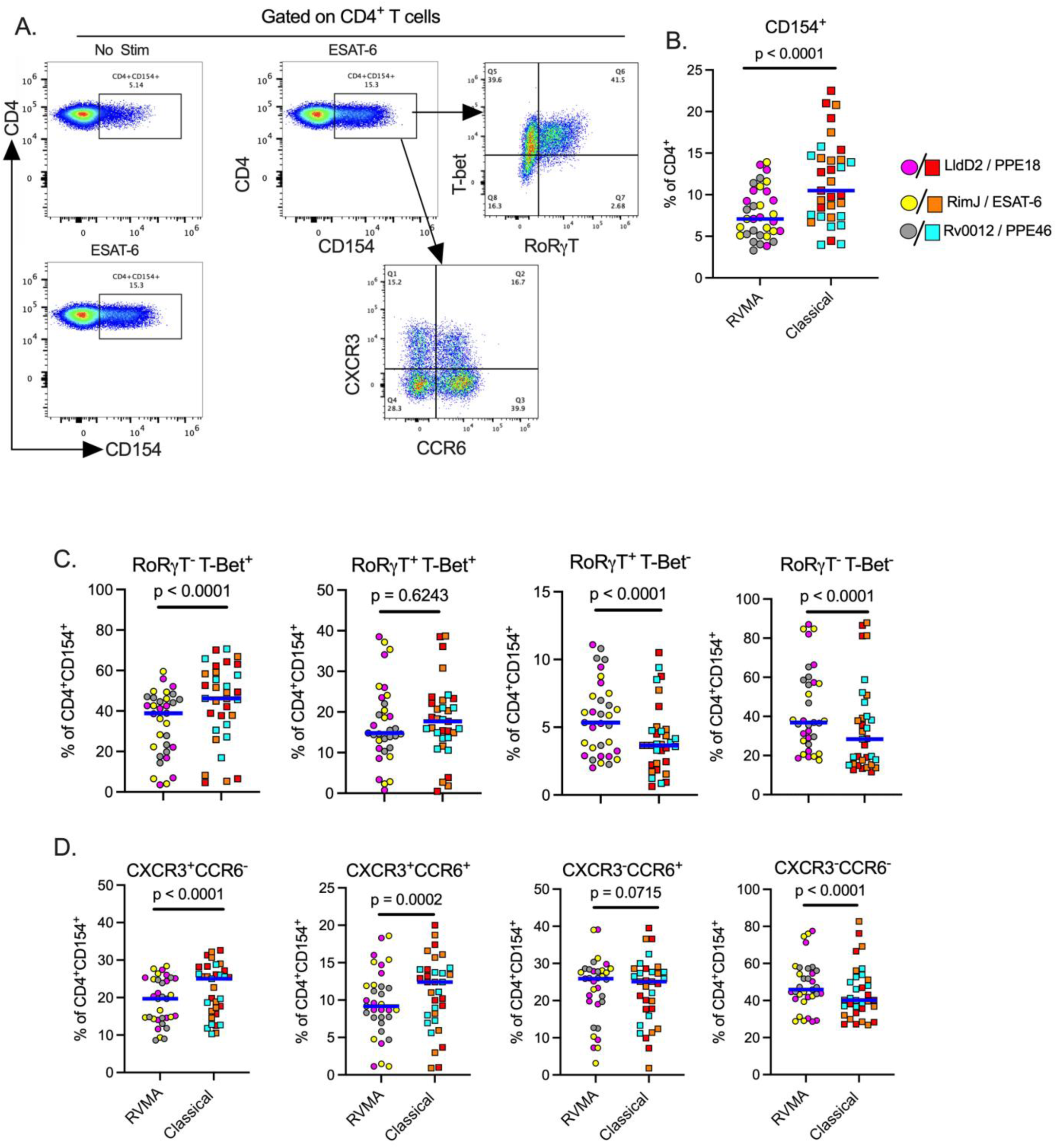
Antigen activated cells express markers of Th17 and Th1 cells upon stimulation with RVMA and classical antigens respectively. Cryopreserved PBMCs from participants in cohort 1 were stimulated with antigens for a total of 20 hours in the presence of costimulatory antibodies anti-CD28 and anti-CD49d and anti-CD40 blocking antibody, and antigen activated cells identified by CD154 surface expression on CD4^+^ T cells. (**A**) Representative flow cytometry plots. (**B**) CD154 surface expression on CD4^+^ T cells, CD154^+^ used to identify antigen activated T cells. (**C**) Expression of helper T cell lineage defining transcription factors (RORγT = Th17) and (T-bet = Th1) and (**D**) Expression of chemokine receptors (CCR6 = Th17 marker) and (CXCR3 = Th1 marker) on antigen activated T cells. Statistics: Wilcoxon matched pairs test.

Since certain chemokine receptors are reported to be associated with specific CD4 T cell subsets, we examined the expression of CCR6 (characteristic of Th17 cells) and CXCR3 (characteristic of Th1 cells). This revealed a significantly higher incidence of CXCR3^+^CCR6^-^ (Th1) and CXCR3^+^CCR6^+^ (Th1*) on antigen-activated (CD4^+^CD154^+^) cells responding to classical antigens than to the RVMA (Fig. 3D), consistent with the previously reported *Mtb*-responsive human Th1* responses to *Mtb* peptide pools (Lindestam Arlehamn et al. 2013; Arlehamn et al. 2014; Nathan et al. 2021; Morgan et al. 2021). In contrast, the incidence of CXCR3^-^CCR6^+^ activated (CD4^+^CD154^+^) cells (Fig. 3D) did not differ by antigen class. Notably, CD4^+^CD154^+^ cells activated by RVMA exhibit a higher incidence of being CXCR3^-^CCR6^-^ than are classical antigen-activated CD4^+^ cells, suggesting that at least some RVMA-responsive cells do not adhere to a conventional pattern of chemokine receptor expression.

## Discussion

In two cohorts of close household contacts that have controlled *Mtb* infection after recent TB exposure, we discovered that *Mtb* antigens -Rv0010c, Rv0012, RimJ and LldD2 - characterized by variable T cell epitopes under diversifying evolutionary selection pressure (Comas et al. 2010; Coscolla et al. 2015) predominantly induce Th17 immunity, which is distinct from T cell responses induced by immunodominant classical *Mtb* antigens that are biased towards Th1 differentiation and IFNγ production. The participants in this study had no signs of TB disease at enrollment and none progressed to TB disease in the 6 - 12 months (cohort 1) or 24 months (cohort 2) of follow-up. We found that CD4 T cells that respond to RVMA express the Th17 lineage-defining transcription factor, RORγT and secrete IL-17 upon antigen stimulation. While we found that some RVMA-responsive CD4 T cells express CCR6, the chemokine receptor associated with Th17 cells, they express CCR6 on proportions of CD4 T cells similar to those that respond to classical antigens. In contrast, we found that a high proportion of RVMA-responsive CD4 T cells express neither CCR6 nor CXCR3 (the chemokine receptor associated with Th1 cells).

Th17 cells are increasingly appreciated as contributing to *Mtb* control in humans (Yu et al. 2017; Domingo-Gonzalez et al. 2017; Milano et al. 2016; Shi and Zhang 2015; Ocejo-Vinyals et al. 2013; Scriba et al. 2017; Nathan et al. 2021; Ogongo et al. 2021) and non-human primate models of *Mtb* infection or vaccine responses (Dijkman et al. 2019; Darrah et al. 2023; 2020; Gideon et al. 2022; Shanmugasundaram et al. 2020) but there has been little investigation of the possibility that Th17 cells might preferentially develop in response to distinct *Mtb* antigens. One important study in which *Mtb* antigens were selected for study in human samples based on their induced expression during infection of multiple strains of mice reported several antigens that induced IL-17 production measured by ELISA in the absence of detectable IFNγ (Coppola et al. 2016). Because of different selection criteria, none of those antigens coincide with the antigens we studied here. Together, the findings in that study and the present one emphasize that *Mtb* can induce CD4 T cells with distinct effector functions, depending on the specific *Mtb* antigen.

Unlike other infectious pathogens in which the targets of protective immunity undergo antigenic variation to escape immune recognition through diversifying evolutionary selection (Crawford et al. 2009; Wang et al. 2010; Rimmelzwaan et al. 2009; Croucher et al. 2011; Seifert et al. 1994; Gkeka et al. 2023; Schneider et al. 2023; Chew et al. 2022), T cell epitopes in the commonly-studied immunodominant *Mtb* antigens are highly conserved(Comas et al. 2010; Coscolla et al. 2015). In this study, we characterized T cells from people protected from progressive/active TB that recognize the rare exceptions, that is, antigens with T cell epitopes that exhibit evidence of diversifying evolutionary selection (variable T cell epitopes) (Coscolla et al. 2015), the RVMA. If *Mtb* follows the evolutionary model of other pathogens, then our results suggest that RVMA are antigens whose recognition by host immune responses is especially detrimental to the pathogen, and we found that recognition of RVMA by T cells is common in people who have controlled *Mtb* infection. Further studies will be needed to determine whether recognition of RVMA differs in those that progress to active TB, but those studies will require human cohorts different than the ones studied here.

It has been widely believed that protective immunity to *Mtb* must involve mechanisms (via cytokines or cytotoxicity) that are directed at *Mtb*-infected cells (usually macrophages). However, our results indicate that T cells that recognize RVMA predominantly express IL-17, which is not thought to directly modulate macrophage microbicidal mechanisms. The mechanisms whereby IL-17 contributes to immunity to *Mtb* have not been fully defined. IL-17 is known to induce expression of multiple antimicrobial peptides, and to induce IL-6, G-CSF, and specific chemokines that promote production and migration of neutrophils (McGeachy et al. 2019). If IL-17 contributes to immunity to *Mtb* through one of these mechanisms, its role may be predominantly in control of extracellular *Mtb* that have escaped from macrophages. In mouse models of *Mtb* infection, IL-17 has been found to induce expression of the chemokine CXCL13 by nonhematopoietic tissues and to contribute to the formation of T and B cell-enriched cellular aggregates in the lungs that are associated with immune control of *Mtb* (Ardain et al. 2019; Gopal et al. 2014; Gopal et al. 2013; Khader et al. 2011). In another mouse model in which *Mtb* infection is characterized by lung tissue necrosis, IL-17 suppressed HIF-1α and tissue hypoxia and reduced lung inflammation (Domingo-Gonzalez et al. 2017). Since evolutionary forces that contribute to pathogen fitness can impact pathogen transmission as well as within-host pathogen survival, it is possible that the effector mechanisms of RVMA-specific T cells predominantly affect *Mtb* transmission. In addition to inducing antimicrobial peptides and neutrophil-directed chemokines, evidence is emerging that IL-17 contributes to tissue homeostasis and repair (reviewed in (McGeachy et al. 2019). Since inflammatory lung tissue destruction and cavitation contribute to *Mtb* transmission (Urbanowski et al. 2020), our results suggest that T cells that target RVMA and produce IL-17 may have their predominant effects on preventing *Mtb* transmission by countering lung tissue destruction. Since pathogen transmission is an important determinant of evolutionary success, antigenic variation to enable *Mtb* to escape recognition by T cells that produce IL-17 and prevent lung tissue damage may account for the sequence diversity of RVMA.

The studies reported here have several limitations. First, although they involved participants in two distinct cohorts in East Africa, they may not be generalizable to other populations. Second, they do not directly reveal whether RVMA-specific human T cells uniquely contribute to protective immunity to *Mtb*; further studies in cohorts that compare responses to RVMA in those with active versus controlled *Mtb* infection will be needed. Third, they do not establish mechanisms whereby RVMA-responsive T cells that produce IL-17 contribute to immunity to *Mtb*.

In summary, we provide unique evidence that *Mtb* antigens with the rare property of undergoing diversifying selection drive development of CD4 T cells with functional properties distinct from T cells that recognize ‘classical’ secreted *Mtb* antigens. Our results suggest that T cells that recognize RVMA make unique contributions to human immunity to TB and may be beneficial antigens to include in TB vaccines. RVMA may be especially valuable in vaccines designed to amplify immune responses that result from initial infection, to broaden the range of T cell effector functions and further reduce the risk of progression to active TB.

## Acknowledgments

The authors thank the members of the TBRU-ASTRa Study Group: Rafi Ahmed, Lance Waller, Lisa Elon, Andrea Knezevic, Shirin Jabbarzadeh, Hao Wu, Seegar Swanson, Yunyun Chen, Wendy Whatney, Melanie Quezada, Loren Sasser, Ranjna Madan Lala, Tawania Fergus, Toidi Adekambi, Deepak Kaushal, Nadia Golden, Taylor Foreman, Allison Bucsan, Chris Ibegbu, Susanna Contraras Alcantra, Alessandro Sette, Salim Allana, Angela Campbell, Sarita Shah, Susan Ray, James Brust, Jeffrey Collins, Meghan Franczek, Jenna Daniel, Anirudh Rao, Rebecca Goldstein, Madeleine Kabongo, Alawode Oladele, Janet Agaya, Dickson Gethi, Dorine Awilly, Albert Ochieng Okumu, Abraham Aseffa, Medina Hamza, Yonas Abebe, Fisseha Mulate, Mekdelawit Wondiyfraw, Firaol Degaga, Daniel Getachew, Dawit Tayachew Bere, Meaza Zewdu, Daniel Mussa, Bezalam Tesfaye, Selam Jemberu, Azeb Tarekegn, Gebeyehu Assefa, Gutema Jebessa, Zewdu Solomon, Sebsibe Neway, Jemal Hussein, Tsegaye Hailu, Alemayehu Geletu, Edom Girma, Million Legesse, Mitin Wendaferew, Hirut Solomon, Zenebech Assefa, Mahlet Mekuria, Misker Kedir, Eleni Zeleke, Rediet Zerihun, Selam Dechasa, Emebet Haile, Nahom Getachew, Firaol Wagari, Ruth Mekonnen, Samuel Bayu, Melat Gebre-Medhin and Alemayehu Kifle. We are also grateful to all study participants and participating health facilities in Kisumu (Kenya) and Addis Ababa (Ethiopia) for their time, dedication, and willingness to participate in the study.

---

**Supplementary Figure 1:**
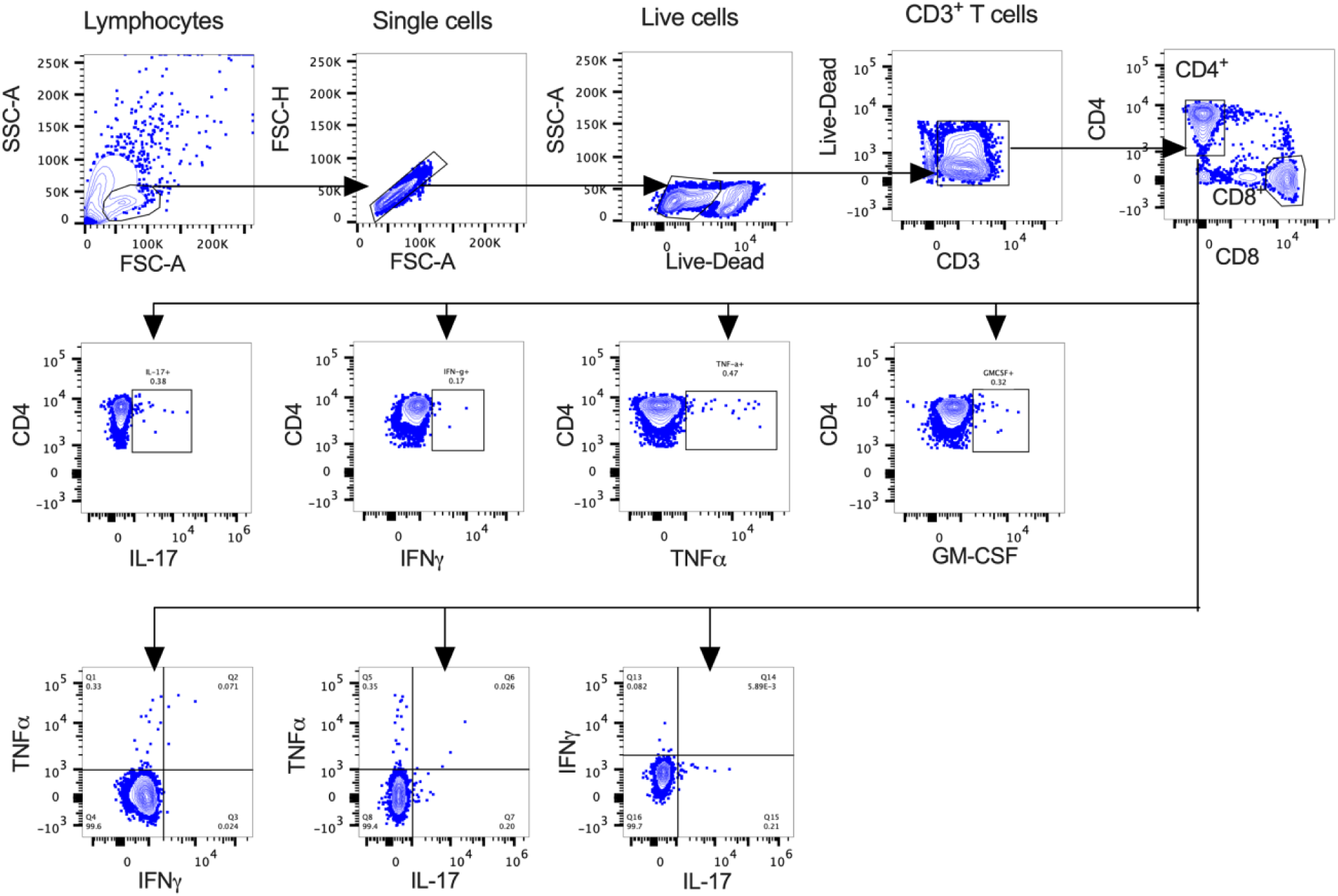
Identification of *Mtb*-specific CD4^+^ T cells. Gating strategy to detect cytokine-producing CD4^+^ T cells after stimulation with distinct *Mtb* antigens. The shown strategy is for unstimulated PBMCs; *Mtb*-specific cytokine magnitude is reported after subtraction of unstimulated background staining.

**Supplementary Figure 2:**
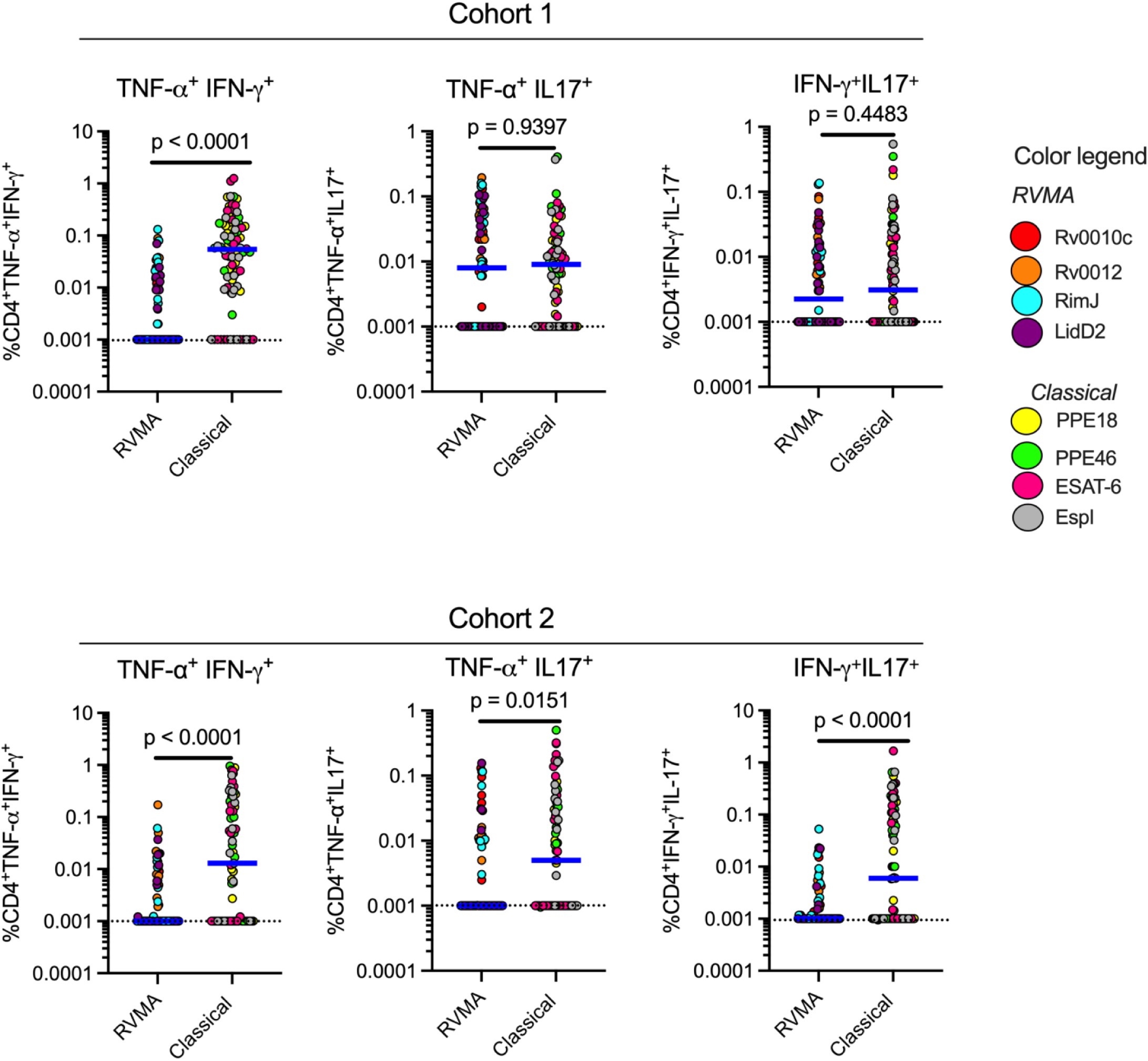
The RVMA induce significantly fewer bifunctional CD4^+^ T cells. Cryopreserved PBMCs from participants in both cohorts were stimulated with distinct antigens (2μg/ml) for a total of 20 hours in the presence of Golgi Stop and Golgi Plug and costimulatory antibodies anti-CD28 and anti-CD49d and dual cytokine production by CD4^+^ T cells determined by intracellular cytokine staining. Each color code is for a distinct antigen as indicated; blue line indicates the median cytokine response. Statistics: Mann-Whitney test.

**Supplemental Table 1.** Demographic and clinical characteristics of study cohorts.

| Characteristic | Cohort 1 | Cohort 2 | p* |
| --- | --- | --- | --- |
| Age, Median (Interquartile range), Y | 31.5 (20, 49.5) | 32.9 (26.7, 38.6) | 0.6613 |
| Sex, n (%) |  |  |  |
| Male | 15 (42%) | 24 (56%) | p = 0.2611,<br>Fisher's exact<br>test) |
| Female | 21 (58%) | 19 (44%) |  |
| BMI; Median (Interquartile range) | 21.95 (20.18, 26.58) | 21.80 (19.8, 24.7) | 0.8852 |
| HbA1c, %; Median (Interquartile range) | 5.5 (5.2, 5.7) | 5.3 (5, 6.1) | 0.7467 |
| QFT Results, IU/mL; Median (Interquartile range) |  |  |  |
| TB antigen minus Nil | 9.03 (1.95, 10) | 5.56 (2.2, 8.23) | **0.0183** |
| Mitogen minus Nil | 5.19 (2.09, 9.46) | 8.78 (7.75, 9.72) | **0.0011** |
| *p = Absolute p, Mann-Whitney unless otherwise specified. |  |  |  |

**Supplemental Table 2.**
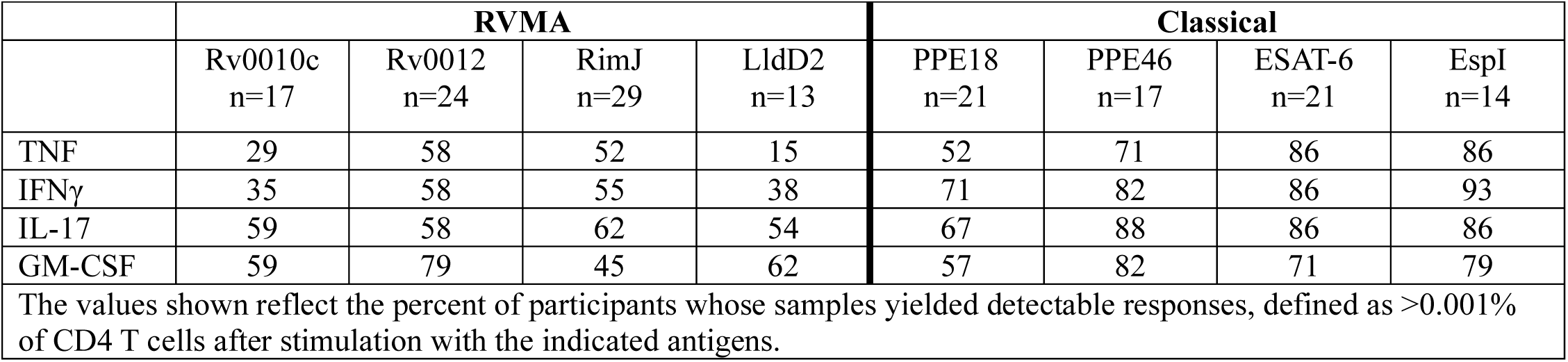
Cohort 2: Frequencies of CD4 T cell cytokine responses, by individual antigens.

**Supplemental Table 3.**
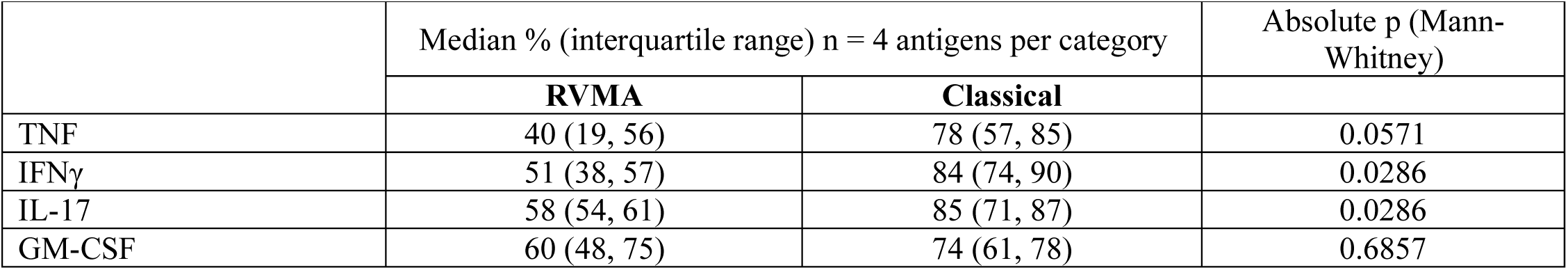
Cohort 2: Frequencies of responders: individual cytokines vs antigen class.

|  | Median % (interquartile range) n = 4 antigens per category |  | Absolute p (Mann-Whitney) |
| --- | --- | --- | --- |
|  | RVMA | Classical |  |
| TNF | 40 (19, 56) | 78 (57, 85) | 0.0571 |
| IFNγ | 51 (38, 57) | 84 (74, 90) | 0.0286 |
| IL-17 | 58 (54, 61) | 85 (71, 87) | 0.0286 |
| GM-CSF | 60 (48, 75) | 74 (61, 78) | 0.6857 |

**Supplemental Table 4.** Cohort 2: Magnitudes of individual cytokine responses (% of CD4 T cells) vs antigen class.

|  | Median % (interquartile range) n = 4 antigens per category |  | Absolute p (Mann-Whitney) |
| --- | --- | --- | --- |
|  | RVMA | Classical |  |
| TNF | 0.001 (0.001, 0.12) | 0.35 (0.001, 0.8) | <0.0001 |
| IFNγ | 0.001 (0.001, 0.06) | 0.3 (0.05, 0.72) | <0.0001 |
| IL-17 | 0.04 (0.001, 0.24) | 0.29 (0.09, 0.66) | <0.0001 |
| GM-CSF | 0.03 (0.001, 0.22) | 0.13 (0.001, 0.32) | 0.0662 |
| Values shown for each cytokine and each antigen are median % of all CD4 T cells that express the specified cytokine. For statistical analyses, assays that yielded undetectable levels of the stated cytokine were assigned a value of 0.001%. |  |  |  |

**Supplemental Table 5.** Cohort 2: IL-17 vs IFNγ responses to individual RVMA.

|  | Response Frequencies (% of participants with detectable cytokine<sup>+</sup> CD4 T cells) |  | Response Magnitudes (% of CD4 T cells that are cytokine<sup>+</sup>) Median (interquartile range) |  |  |
| --- | --- | --- | --- | --- | --- |
|  | **IL-17** | **IFNγ** | **IL-17** | **IFNγ** | **p\*** |
| Rv0010c | 58 | 35 | 0.02 (0.001, 0.115) | 0.001 (0.001, 0.047) | 0.2437 |
| Rv0012 | 58 | 58 | 0.07 (0.001, 0.385) | 0.012 (0.001, 0.0775) | 0.0054 |
| RimJ | 62 | 55 | 0.06 (0.001, 0.32) | 0.01 (0.001, 0.087) | 0.2157 |
| LldD2 | 63 | 46 | 0.03 (0.001, 0.255) | 0.001 (0.001,0.0305) | 0.0391 |
| *p values for the comparison of IL-17 vs IFNγ magnitudes (Wilcoxon matched pairs) | | | | | |

## References

Alwis, Ruklanthi de, Katherine L. Williams, Michael A. Schmid, Chih-Yun Lai, Bhumi Patel, Scott A. Smith, James E. Crowe, Wei-Kung Wang, Eva Harris, and Aravinda M. de Silva. 2014. “Dengue Viruses Are Enhanced by Distinct Populations of Serotype Cross-Reactive Antibodies in Human Immune Sera.” PLoS Pathogens 10 (10): e1004386. 10.1371/journal.ppat.1004386.

Ardain, Amanda, Racquel Domingo-Gonzalez, Shibali Das, Samuel W. Kazer, Nicole C. Howard, Alveera Singh, Mushtaq Ahmed, et al. 2019. “Group 3 Innate Lymphoid Cells Mediate Early Protective Immunity against Tuberculosis.” *Nature*, June. 10.1038/s41586-019-1276-2.

Arlehamn, Cecilia Lindestam, Gregory Seumois, Anna Gerasimova, Charlie Huang, Zheng Fu, Xiaojing Yue, Alessandro Sette, Pandurangan Vijayanand, and Bjoern Peters. 2014. “Transcriptional Profile of TB Antigen-Specific T Cells Reveals Novel Multifunctional Features.” *Journal of Immunology (Baltimore*, Md*. :* 1950) 193 (6): 2931–40. 10.4049/jimmunol.1401151.

Barham, Morgan S., Wendy E. Whatney, Jeremiah Khayumbi, Joshua Ongalo, Loren E. Sasser, Angela Campbell, Meghan Franczek, et al. 2020. “Activation-Induced Marker Expression Identifies Mycobacterium Tuberculosis-Specific CD4 T Cells in a Cytokine-Independent Manner in HIV-Infected Individuals with Latent Tuberculosis.” ImmunoHorizons 4 (10): 573–84. 10.4049/immunohorizons.2000051.

Barnes, P. F., A. B. Bloch, P. T. Davidson, and D. E. Snider. 1991. “Tuberculosis in Patients with Human Immunodeficiency Virus Infection.” The New England Journal of Medicine 324 (23): 1644–50. 10.1056/NEJM199106063242307.

Chew, Marvin, Weijian Ye, Radoslaw Igor Omelianczyk, Charisse Flerida Pasaje, Regina Hoo, Qingfeng Chen, Jacquin C. Niles, Jianzhu Chen, and Peter Preiser. 2022. “Selective Expression of Variant Surface Antigens Enables Plasmodium Falciparum to Evade Immune Clearance in Vivo.” Nature Communications 13 (1): 4067. 10.1038/s41467-022-31741-2.

Cole, S. T., R. Brosch, J. Parkhill, T. Garnier, C. Churcher, D. Harris, S. V. Gordon, et al. 1998. “Deciphering the Biology of Mycobacterium Tuberculosis from the Complete Genome Sequence.” Nature 393 (6685): 537–44. 10.1038/31159.

Comas, Iñaki, Jaidip Chakravartti, Peter M. Small, James Galagan, Stefan Niemann, Kristin Kremer, Joel D. Ernst, and Sebastien Gagneux. 2010. “Human T Cell Epitopes of Mycobacterium Tuberculosis Are Evolutionarily Hyperconserved.” Nature Genetics 42 (6): 498–503. 10.1038/ng.590.

Coppola, Mariateresa, Krista E. van Meijgaarden, Kees L. M. C. Franken, Susanna Commandeur, Gregory Dolganov, Igor Kramnik, Gary K. Schoolnik, et al. 2016. “New Genome-Wide Algorithm Identifies Novel In-Vivo Expressed Mycobacterium Tuberculosis Antigens Inducing Human T-Cell Responses with Classical and Unconventional Cytokine Profiles.” Scientific Reports 6: 37793. 10.1038/srep37793.

Coscolla, Mireia, Richard Copin, Jayne Sutherland, Florian Gehre, Bouke de Jong, Olumuiya Owolabi, Georgetta Mbayo, Federica Giardina, Joel D. Ernst, and Sebastien Gagneux. 2015. “M. Tuberculosis T Cell Epitope Analysis Reveals Paucity of Antigenic Variation and Identifies Rare Variable TB Antigens.” Cell Host & Microbe 18 (5): 538–48. 10.1016/j.chom.2015.10.008.

Crawford, Hayley, Wendy Lumm, Alasdair Leslie, Malinda Schaefer, Debrah Boeras, Julia G. Prado, Jianming Tang, et al. 2009. “Evolution of HLA-B*5703 HIV-1 Escape Mutations in HLA-B*5703-Positive Individuals and Their Transmission Recipients.” The Journal of Experimental Medicine 206 (4): 909–21. 10.1084/jem.20081984.

Croucher, Nicholas J., Simon R. Harris, Christophe Fraser, Michael A. Quail, John Burton, Mark van der Linden, Lesley McGee, et al. 2011. “Rapid Pneumococcal Evolution in Response to Clinical Interventions.” Science (New York, N.Y.) 331 (6016): 430–34. 10.1126/science.1198545.

Dan, Jennifer M., Cecilia S. Lindestam Arlehamn, Daniela Weiskopf, Ricardo da Silva Antunes, Colin Havenar-Daughton, Samantha M. Reiss, Matthew Brigger, Marcella Bothwell, Alessandro Sette, and Shane Crotty. 2016. “A Cytokine-Independent Approach To Identify Antigen-Specific Human Germinal Center T Follicular Helper Cells and Rare Antigen-Specific CD4+ T Cells in Blood.” Journal of Immunology (Baltimore, Md.: 1950) 197 (3): 983–93. 10.4049/jimmunol.1600318.

Darrah, Patricia A., Joseph J. Zeppa, Pauline Maiello, Joshua A. Hackney, Marc H. Wadsworth, Travis K. Hughes, Supriya Pokkali, et al. 2020. “Prevention of Tuberculosis in Macaques after Intravenous BCG Immunization.” Nature 577 (7788): 95–102. 10.1038/s41586-019-1817-8.

Darrah, Patricia A., Joseph J. Zeppa, Chuangqi Wang, Edward B. Irvine, Allison N. Bucsan, Mark A. Rodgers, Supriya Pokkali, et al. 2023. “Airway T Cells Are a Correlate of i.v. Bacille Calmette-Guerin-Mediated Protection against Tuberculosis in Rhesus Macaques.” Cell Host & Microbe 0 (0). 10.1016/j.chom.2023.05.006.

Delgado, Maria Florencia, Silvina Coviello, A. Clara Monsalvo, Guillermina A. Melendi, Johanna Zea Hernandez, Juan P. Batalle, Leandro Diaz, et al. 2009. “Lack of Antibody Affinity Maturation Due to Poor Toll-like Receptor Stimulation Leads to Enhanced Respiratory Syncytial Virus Disease.” Nature Medicine 15 (1): 34–41. 10.1038/nm.1894.

Dijkman, Karin, Claudia C. Sombroek, Richard A. W. Vervenne, Sam O. Hofman, Charelle Boot, Edmond J. Remarque, Clemens H. M. Kocken, et al. 2019. “Prevention of Tuberculosis Infection and Disease by Local BCG in Repeatedly Exposed Rhesus Macaques.” Nature Medicine 25 (2): 255–62. 10.1038/s41591-018-0319-9.

Domingo-Gonzalez, Racquel, Shibali Das, Kristin L. Griffiths, Mushtaq Ahmed, Monika Bambouskova, Radha Gopal, Suhas Gondi, et al. 2017. “Interleukin-17 Limits Hypoxia-Inducible Factor 1α and Development of Hypoxic Granulomas during Tuberculosis.” JCI Insight 2 (19). 10.1172/jci.insight.92973.

Flory, C. M., R. D. Hubbard, and F. M. Collins. 1992. “Effects of in Vivo T Lymphocyte Subset Depletion on Mycobacterial Infections in Mice.” Journal of Leukocyte Biology 51 (3): 225–29.

Gideon, Hannah P., Travis K. Hughes, Constantine N. Tzouanas, Marc H. Wadsworth, Ang Andy Tu, Todd M. Gierahn, Joshua M. Peters, et al. 2022. “Multimodal Profiling of Lung Granulomas in Macaques Reveals Cellular Correlates of Tuberculosis Control.” Immunity 55 (5): 827–846.e10. 10.1016/j.immuni.2022.04.004.

Gkeka, Anastasia, Francisco Aresta-Branco, Gianna Triller, Evi P. Vlachou, Monique van Straaten, Mirjana Lilic, Paul Dominic B. Olinares, et al. 2023. “Immunodominant Surface Epitopes Power Immune Evasion in the African Trypanosome.” Cell Reports 42 (3): 112262. 10.1016/j.celrep.2023.112262.

Goldschneider, I., E. C. Gotschlich, and M. S. Artenstein. 1969. “Human Immunity to the Meningococcus. I. The Role of Humoral Antibodies.” The Journal of Experimental Medicine 129 (6): 1307–26. 10.1084/jem.129.6.1307.

Gopal, R., J. Rangel-Moreno, S. Slight, Y. Lin, H. F. Nawar, B. A. Fallert Junecko, T. A. Reinhart, et al. 2013. “Interleukin-17-Dependent CXCL13 Mediates Mucosal Vaccine-Induced Immunity against Tuberculosis.” Mucosal Immunology 6 (5): 972–84. 10.1038/mi.2012.135.

Gopal, Radha, Leticia Monin, Samantha Slight, Uzodinma Uche, Emmeline Blanchard, Beth A. Fallert Junecko, Rosalio Ramos-Payan, et al. 2014. “Unexpected Role for IL-17 in Protective Immunity against Hypervirulent Mycobacterium Tuberculosis HN878 Infection.” PLoS Pathogens 10 (5): e1004099. 10.1371/journal.ppat.1004099.

Grifoni, Alba, Daniela Weiskopf, Sydney I. Ramirez, Jose Mateus, Jennifer M. Dan, Carolyn Rydyznski Moderbacher, Stephen A. Rawlings, et al. 2020. “Targets of T Cell Responses to SARS-CoV-2 Coronavirus in Humans with COVID-19 Disease and Unexposed Individuals.” Cell, May. 10.1016/j.cell.2020.05.015.

Hoyos, David, Roberta Zappasodi, Isabell Schulze, Zachary Sethna, Kelvin César de Andrade, Dean F. Bajorin, Chaitanya Bandlamudi, et al. 2022. “Fundamental Immune-Oncogenicity Trade-Offs Define Driver Mutation Fitness.” Nature 606 (7912): 172–79. 10.1038/s41586-022-04696-z.

Katzelnick, Leah C., Lionel Gresh, M. Elizabeth Halloran, Juan Carlos Mercado, Guillermina Kuan, Aubree Gordon, Angel Balmaseda, and Eva Harris. 2017. “Antibody-Dependent Enhancement of Severe Dengue Disease in Humans.” *Science (New York*, N.Y.) 358 (6365): 929–32. 10.1126/science.aan6836.

Khader, Shabaana A., Lokesh Guglani, Javier Rangel-Moreno, Radha Gopal, Beth A. Fallert Junecko, Jeffrey J. Fountain, Cynthia Martino, et al. 2011. “IL-23 Is Required for Long-Term Control of Mycobacterium Tuberculosis and B Cell Follicle Formation in the Infected Lung.” *Journal of Immunology (Baltimore*, Md.: 1950) 187 (10): 5402–7. 10.4049/jimmunol.1101377.

Kiepiela, Photini, Kholiswa Ngumbela, Christina Thobakgale, Dhanwanthie Ramduth, Isobella Honeyborne, Eshia Moodley, Shabashini Reddy, et al. 2007. “CD8+ T-Cell Responses to Different HIV Proteins Have Discordant Associations with Viral Load.” Nature Medicine 13 (1): 46–53. 10.1038/nm1520.

Lawn, Stephen D., Landon Myer, David Edwards, Linda-Gail Bekker, and Robin Wood. 2009. “Short-Term and Long-Term Risk of Tuberculosis Associated with CD4 Cell Recovery during Antiretroviral Therapy in South Africa.” *AIDS (London*, England) 23 (13): 1717–25. 10.1097/QAD.0b013e32832d3b6d.

Leveton, C., S. Barnass, B. Champion, S. Lucas, B. De Souza, M. Nicol, D. Banerjee, and G. Rook. 1989. “T-Cell-Mediated Protection of Mice against Virulent Mycobacterium Tuberculosis.” Infection and Immunity 57 (2): 390–95. 10.1128/iai.57.2.390-395.1989.

Lindestam Arlehamn, Cecilia S., Anna Gerasimova, Federico Mele, Ryan Henderson, Justine Swann, Jason A. Greenbaum, Yohan Kim, et al. 2013. “Memory T Cells in Latent Mycobacterium Tuberculosis Infection Are Directed against Three Antigenic Islands and Largely Contained in a CXCR3+CCR6+ Th1 Subset.” PLoS Pathogens 9 (1): e1003130. 10.1371/journal.ppat.1003130.

Marty Pyke, Rachel, Wesley Kurt Thompson, Rany M. Salem, Joan Font-Burgada, Maurizio Zanetti, and Hannah Carter. 2018. “Evolutionary Pressure against MHC Class II Binding Cancer Mutations.” Cell 175 (2): 416–428.e13. 10.1016/j.cell.2018.08.048.

Marty, Rachel, Saghar Kaabinejadian, David Rossell, Michael J. Slifker, Joris van de Haar, Hatice Billur Engin, Nicola de Prisco, et al. 2017. “MHC-I Genotype Restricts the Oncogenic Mutational Landscape.” Cell 171 (6): 1272–1283.e15. 10.1016/j.cell.2017.09.050.

Matsushita, Hirokazu, Matthew D. Vesely, Daniel C. Koboldt, Charles G. Rickert, Ravindra Uppaluri, Vincent J. Magrini, Cora D. Arthur, et al. 2012. “Cancer Exome Analysis Reveals a T-Cell-Dependent Mechanism of Cancer Immunoediting.” Nature 482 (7385): 400–404. 10.1038/nature10755.

McGeachy, Mandy J., Daniel J. Cua, and Sarah L. Gaffen. 2019. “The IL-17 Family of Cytokines in Health and Disease.” Immunity 50 (4): 892–906. 10.1016/j.immuni.2019.03.021.

Milano, Mariana, Milton Ozório Moraes, Rodrigo Rodenbusch, Caroline Xavier Carvalho, Melaine Delcroix, Gabriel Mousquer, Lucas Laux da Costa, Gisela Unis, Elis Regina Dalla Costa, and Maria Lucia Rosa Rossetti. 2016. “Single Nucleotide Polymorphisms in IL17A and IL6 Are Associated with Decreased Risk for Pulmonary Tuberculosis in Southern Brazilian Population.” PloS One 11 (2): e0147814. 10.1371/journal.pone.0147814.

Mogues, T., M. E. Goodrich, L. Ryan, R. LaCourse, and R. J. North. 2001. “The Relative Importance of T Cell Subsets in Immunity and Immunopathology of Airborne Mycobacterium Tuberculosis Infection in Mice.” The Journal of Experimental Medicine 193 (3): 271–80.

Morgan, Jeffrey, Kaylin Muskat, Rashmi Tippalagama, Alessandro Sette, Julie Burel, and Cecilia S. Lindestam Arlehamn. 2021. “Classical CD4 T Cells as the Cornerstone of Antimycobacterial Immunity.” Immunological Reviews 301 (1): 10–29. 10.1111/imr.12963.

Musvosvi, Munyaradzi, Huang Huang, Chunlin Wang, Qiong Xia, Virginie Rozot, Akshaya Krishnan, Peter Acs, et al. 2023. “T Cell Receptor Repertoires Associated with Control and Disease Progression Following Mycobacterium Tuberculosis Infection.” Nature Medicine 29 (1): 258–69. 10.1038/s41591-022-02110-9.

Nathan, Aparna, Jessica I. Beynor, Yuriy Baglaenko, Sara Suliman, Kazuyoshi Ishigaki, Samira Asgari, Chuan-Chin Huang, et al. 2021. “Multimodally Profiling Memory T Cells from a Tuberculosis Cohort Identifies Cell State Associations with Demographics, Environment and Disease.” Nature Immunology 22 (6): 781–93. 10.1038/s41590-021-00933-1.

Ocejo-Vinyals, Javier Gonzalo, Elena Puente de Mateo, María Ángeles Hoz, José Luis Arroyo, Ramón Agüero, Fernado Ausín, and M. Carmen Fariñas. 2013. “The IL-17 G-152A Single Nucleotide Polymorphism Is Associated with Pulmonary Tuberculosis in Northern Spain.” Cytokine 64 (1): 58–61. 10.1016/j.cyto.2013.05.022.

Ogongo, Paul, Liku B. Tezera, Amanda Ardain, Shepherd Nhamoyebonde, Duran Ramsuran, Alveera Singh, Abigail Ng’oepe, et al. 2021. “Tissue-Resident-like CD4+ T Cells Secreting IL-17 Control Mycobacterium Tuberculosis in the Human Lung.” The Journal of Clinical Investigation 131 (10): 142014. 10.1172/JCI142014.

Polack, Fernando P., Michael N. Teng, Peter L. Collins, Gregory A. Prince, Marcus Exner, Heinz Regele, Dario D. Lirman, et al. 2002. “A Role for Immune Complexes in Enhanced Respiratory Syncytial Virus Disease.” The Journal of Experimental Medicine 196 (6): 859–65. 10.1084/jem.20020781.

Ranasinghe, Srinika, Michael Flanders, Sam Cutler, Damien Z. Soghoian, Musie Ghebremichael, Isaiah Davis, Madelene Lindqvist, et al. 2012. “HIV-Specific CD4 T Cell Responses to Different Viral Proteins Have Discordant Associations with Viral Load and Clinical Outcome.” Journal of Virology 86 (1): 277–83. 10.1128/JVI.05577-11.

Rimmelzwaan, Guus F., Joost H. C. M. Kreijtz, Rogier Bodewes, Ron A. M. Fouchier, and Albert D. M. E. Osterhaus. 2009. “Influenza Virus CTL Epitopes, Remarkably Conserved and Remarkably Variable.” Vaccine 27 (45): 6363–65. 10.1016/j.vaccine.2009.01.016.

Scanga, C. A., V. P. Mohan, K. Yu, H. Joseph, K. Tanaka, J. Chan, and J. L. Flynn. 2000. “Depletion of CD4(+) T Cells Causes Reactivation of Murine Persistent Tuberculosis despite Continued Expression of Interferon Gamma and Nitric Oxide Synthase 2.” The Journal of Experimental Medicine 192 (3): 347–58. 10.1084/jem.192.3.347.

Schneider, Victoria M., Joseph E. Visone, Chantal T. Harris, Francesca Florini, Evi Hadjimichael, Xu Zhang, Mackensie R. Gross, et al. 2023. “The Human Malaria Parasite Plasmodium Falciparum Can Sense Environmental Changes and Respond by Antigenic Switching.” Proceedings of the National Academy of Sciences of the United States of America 120 (17): e2302152120. 10.1073/pnas.2302152120.

Scriba, Thomas J., Adam Penn-Nicholson, Smitha Shankar, Tom Hraha, Ethan G. Thompson, David Sterling, Elisa Nemes, et al. 2017. “Sequential Inflammatory Processes Define Human Progression from M. Tuberculosis Infection to Tuberculosis Disease.” PLoS Pathogens 13 (11): e1006687. 10.1371/journal.ppat.1006687.

Seifert, H. S., C. J. Wright, A. E. Jerse, M. S. Cohen, and J. G. Cannon. 1994. “Multiple Gonococcal Pilin Antigenic Variants Are Produced during Experimental Human Infections.” The Journal of Clinical Investigation 93 (6): 2744–49. 10.1172/JCI117290.

Shanmugasundaram, Uma, Allison N. Bucsan, Shashank R. Ganatra, Chris Ibegbu, Melanie Quezada, Robert V. Blair, Xavier Alvarez, Vijayakumar Velu, Deepak Kaushal, and Jyothi Rengarajan. 2020. “Pulmonary Mycobacterium Tuberculosis Control Associates with CXCR3- and CCR6-Expressing Antigen-Specific Th1 and Th17 Cell Recruitment.” JCI Insight 5 (14): 137858. 10.1172/jci.insight.137858.

Shi, G.-C., and L.-G. Zhang. 2015. “Influence of Interleukin-17 Gene Polymorphisms on the Development of Pulmonary Tuberculosis.” Genetics and Molecular Research: GMR 14 (3): 8526–31. 10.4238/2015.July.28.22.

Sonnenberg, Pam, Judith R. Glynn, Katherine Fielding, Jill Murray, Peter Godfrey-Faussett, and Stuart Shearer. 2005. “How Soon after Infection with HIV Does the Risk of Tuberculosis Start to Increase? A Retrospective Cohort Study in South African Gold Miners.” The Journal of Infectious Diseases 191 (2): 150–58. 10.1086/426827.

Urbanowski, Michael E., Alvaro A. Ordonez, Camilo A. Ruiz-Bedoya, Sanjay K. Jain, and William R. Bishai. 2020. “Cavitary Tuberculosis: The Gateway of Disease Transmission.” The Lancet. Infectious Diseases 20 (6): e117–28. 10.1016/S1473-3099(20)30148-1.

Wang, Shuping, Rico Buchli, Jennifer Schiller, Jianen Gao, Rodney S. VanGundy, William H. Hildebrand, and David D. Eckels. 2010. “Natural Epitope Variants of the Hepatitis C Virus Impair Cytotoxic T Lymphocyte Activity.” World Journal of Gastroenterology 16 (16): 1953–69. 10.3748/wjg.v16.i16.1953.

Whatney, Wendy E., Neel R. Gandhi, Cecilia S. Lindestam Arlehamn, Azhar Nizam, Hao Wu, Melanie J. Quezada, Angela Campbell, et al. 2018. “A High Throughput Whole Blood Assay for Analysis of Multiple Antigen-Specific T Cell Responses in Human Mycobacterium Tuberculosis Infection.” *Journal of Immunology (Baltimore*, Md.: 1950) 200 (8): 3008–19. 10.4049/jimmunol.1701737.

Yu, Z.-G., B.-Z. Wang, J. Li, Z.-L. Ding, and K. Wang. 2017. “Association between Interleukin-17 Genetic Polymorphisms and Tuberculosis Susceptibility: An Updated Meta-Analysis.” The International Journal of Tuberculosis and Lung Disease: The Official Journal of the International Union Against Tuberculosis and Lung Disease 21 (12): 1307–13. 10.5588/ijtld.17.0345.

